# Both stimulus-specific and configurational features of multiple visual stimuli shape the spatial ventriloquism effect

**DOI:** 10.1101/2023.05.02.539018

**Authors:** Christoph Kayser, Nienke Debats, Herbert Heuer

## Abstract

Studies on multisensory perception often focus on simplistic conditions in which one single stimulus is presented per modality. Yet, in everyday life we usually encounter multiple signals per modality. To understand how multiple signals within and across the senses are combined we extended the classical audio-visual spatial ventriloquism paradigm to combine two visual stimuli with one sound. The individual visual stimuli presented in the same trial differed in their relative timing and spatial offsets to the sound, allowing us to contrast their individual and combined influence on sound localization judgements. We find that the ventriloquism bias is not dominated by a single visual stimulus but rather is shaped by the collective multisensory evidence. In particular, the contribution of an individual visual stimulus to the ventriloquism bias depends not only on its own relative spatio-temporal alignment to the sound but also the spatio-temporal alignment of the other visual stimulus. We propose that this pattern of multi-stimulus multisensory integration reflects the evolution of evidence for sensory causal relations during individual trials, calling for the need to extend established models of multisensory causal inference to more naturalistic conditions. Our data also suggest that this pattern of multisensory interactions extends to the ventriloquism aftereffect, a bias in sound localization observed in unisensory judgements following a multisensory stimulus.

## Introduction

Multisensory perception involves the binding of stimuli across sensory modalities. In real life, we usually encounter multiple cues in each modality that can belong to the same or distinct objects. For example, while hearing the hiss of a snake we may spot movements at multiple locations on the ground, only one of which gives cues about the snake’s location. This raises the question of how the presence of more than one stimulus per modality influences multisensory perception. While it is known that multisensory binding is generally shaped by the spatio-temporal relation of the stimuli, their semantic attributes, their task relevance and the attentional focus (Talsma *et al*., 2010; Noppeney, 2021; Shams & Beierholm, 2022), still little is known about how multiple signals from one modality are combined with one signal from another.

We here address this question using the spatial ventriloquism paradigm as a model (Howard & Templeton, 1966; Bertelson *et al*., 2000; Bruns, 2019). Across four experiments we asked how two visual signals are combined with one sound to shape judgements about the sound’s location. During the key trials in these experiments one sound could be accompanied by one or two visual stimuli that differed in their relative timing and spatial offset to the sound. Conceptually, one could envisage different patterns of multisensory interactions to emerge in this setting. One possibility is that one visual signal dominates (or captures) the binding process, for example the one most spatially proximal or synchronous with the sound (Soto-Faraco *et al*., 2002; Alink *et al*., 2008; Van der Burg *et al*., 2008a; Stekelenburg & Vroomen, 2009; Bruns & Roder, 2010). While this is arguably somewhat unlikely, sensory capture has remained a prominent framework in multisensory perception (Colavita, 1974; Spence, 2009; Chen & Spence, 2017). Alternatively, all stimuli could contribute to the sound localization judgements in a graded manner (Landy *et al*., 1995; Ernst & Banks, 2002; Ernst & Bulthoff, 2004; Stein & Stanford, 2008; Angelaki *et al*., 2009; Ohshiro *et al*., 2011). Hereby, the relative influence of each visual stimulus could either depend only on its own spatio-temporal distance to the sound, or alternatively, may also depend on the overall context provided by the presence of other visual signals (Bertelson *et al*., 2000). In the former case the multisensory bias observed with two visual stimuli would reflect the superposition of the influences of the individual visual stimuli. In the latter case it would reflect more elaborate multisensory computations that shape sound localization based on the collective sensory evidence, possibly in a non-linear manner as incorporated in models of multisensory causal inference (Kording *et al*., 2007; McGovern *et al*., 2016; Noppeney, 2021; Shams & Beierholm, 2022). We here compared these alternatives based on dedicated experimental manipulations and the analysis of participants’ response biases in conditions involving two independent visual stimuli in the same trial.

To date, few studies have investigated multisensory perception under conditions where two stimuli in one modality are presented together with one in another. Using a visuo-motor paradigm, we have tested how the presence of two visual cues affects judgements about the proprioceptively-felt hand position (Debats *et al*., 2021). In that study participants made center-out hand movements and were presented with two spatially- and temporally-offset visual signals that provided discrepant evidence about their movement direction. Participants’ judgements about movement endpoints were shaped by both visual signals, pointing to the weighted influence of multiple signals within and across modalities. In the audio-visual domain, previous work showed that the interaction between multiple visual stimuli and one sound is shaped by semantic congruency (Kanaya & Yokosawa, 2011; Shavit-Cohen & Zion Golumbic, 2019), temporal synchronization (Lewkowicz *et al*., 2021) and prior exposure to these stimuli (Tong *et al*., 2020). Furthermore, in a spatial ventriloquism experiment with two visual stimuli flanking one sound at equal spatial distances, the more salient visual signal dominated the binding process (Bertelson *et al*., 2000). However, when both visual signals were equally salient their effects cancelled, supporting the idea that multiple signals within a modality collectively shape multisensory binding.

To provide a comprehensive assessment of multisensory perception we probed two perceptual biases that are typically studied in this experimental paradigm: the ventriloquism effect, describing how visual and acoustic evidence in a given audio-visual trial is combined to judge the location of these stimuli (Bertelson & Radeau, 1981; Rohe & Noppeney, 2015), and the immediate (trial-wise) ventriloquism aftereffect. The latter reflects a bias in the judgement of unisensory auditory stimuli that are presented following a multisensory trial and is considered an instance of crossmodal adaptation or learning (Radeau & Bertelson, 1974; Recanzone, 1998; Bruns *et al*., 2011; Park & Kayser, 2019; 2021).

The outline of the study is as follows. In a first experiment we probed the temporal pattern of multisensory binding by manipulating the stimulus onset asynchrony (SOA) between one visual stimulus and a sound. This revealed a comparable influence of visual stimuli that precede or follow the sound and allowed us to establish a suitable SOA magnitude for the subsequent experiments. In three experiments we then probed how the presence of one or two visual stimuli contributes to sound localization judgements. Two experiments were designed such that two visual stimuli (presented at distinct SOAs) featured spatial offsets relative to the sound of same magnitude but opposing directions. Hence, if the two visual stimuli would exert a comparable influence on sound localization judgements the collective response bias should vanish. Alternatively, if one visual stimulus would dominate the binding process the collective bias should be comparable to that observed in a condition where only the dominant visual stimulus is present. Finally, in a last experiment we manipulated the relative spatial discrepancies of the two visual stimuli independently, so that they could feature discrepancies of same or distinct magnitude and same and distinct directions. This allowed us to verify that results generalize to conditions in which the number of visual stimuli, their timing to the sound, and their spatial arrangement are unpredictable.

## METHODS

### Participants

All studies were approved by the ethics committee of Bielefeld University. Adult volunteers participated after providing informed consent and were compensated for their time. All had self-reported normal vision and hearing and none indicated a history of neurological disorders. The data were collected anonymously and it is possible that some individuals participated in more than one of the experiments described below. Demographical data was not collected. For each experiment we aimed for a sample size of at least 20 participants, following previous studies using similar experimental protocols (Park & Kayser, 2019; Park *et al*., 2021; Park & Kayser, 2022) and recommendations for sample sizes in experimental studies (Simmons *et al*., 2011). All in all, we invited 95 individuals to these experiments. Four of these did not pass the screening test for spatial hearing and were not invited for the main experiments. Of the collected 91 full datasets four were removed as detailed below.

### General experimental setup and paradigm

The general paradigm was based on a single-trial audio-visual localization task in which the individual trials and conditions are designed to probe the ventriloquism effect and the immediate (trial-wise) ventriloquism aftereffect (Wozny & Shams, 2011b; Park & Kayser, 2019; 2021; 2022) (Figure 1). Participants were seated in an echo-free booth (E:Box, Desone) in front of an acoustically transparent screen (Screen International Modigliani, 2×1 m^2^, about 1m from the participants head) with their head supported on a chin rest. The experiments generally comprised audio-visual (AV) trials, auditory (A) trials and visual (V) trials, and participants were asked to localize either the sound (in AV or A trials) or a visual stimulus (in AV or V trials; for details see individual experiments below). The main effects of interest were probed based on AV trials requiring participants to judge the location of the sound. The pure visual trials or visual judgements in AV trials were mostly included to maintain attention divided across modalities. Each trial started with a fixation period (uniform 700 to 1100 ms), followed by the stimulus presentation. The individual stimuli lasted 50 ms but were possibly presented with different stimulus onset asynchronies (SOA). After a post-stimulus interval (uniform 400 to 700 ms) a response cue was presented. This was a horizontal bar along which participants could move a mouse cursor and indicate their judgement by clicking the mouse button. A letter instructed participants about which stimulus they were supposed to localize. Inter-trial intervals varied randomly (uniform 900 to 1200 ms). Participants were asked to maintain fixation during the fixation and stimulus periods but could move their eyes when responding and in inter-trial intervals. Eye movements were not monitored but stimuli had sufficiently short durations to avoid eye movements during stimulus presentation; see (Bosen *et al*., 2017; Park & Kayser, 2019; 2021) for a discussion of a potential role of eye movements in these paradigms.

**Figure 1.**
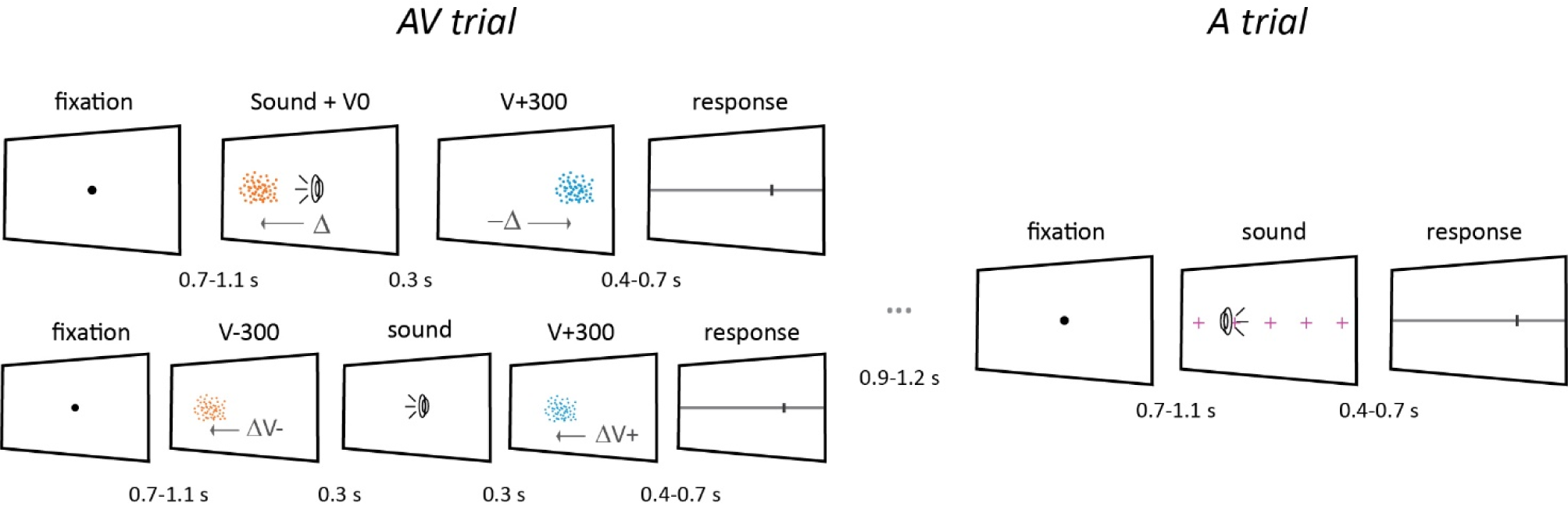
Schematic of experimental trials. The precise design and parameters differed between the four experiments. Each trial consisted of a fixation interval, stimulus presentation and a response interval. Common to all experiments is the audio-visual trial (AV), in which a sound was presented together with one or two visual stimuli. In the AV trials of experiment 1 only one visual stimulus was presented and could appear at one of five stimulus onset asynchronies (SOA: ±600 ms, ±300 ms, and 0 ms) relative to the sound. In experiments 2 to 4 either one or two visual stimuli were presented, as shown on the left. In experiment 2 the visual stimuli could be synchronous (V0) or delayed to the sound (by 300 ms, V+300), in experiments 3 and 4 they could either precede the sound by 300 ms (V-300), be synchronous (V0), or delayed by 300 ms (V+300). In experiments 2 and 3, when two visual stimuli were presented in an AV trial they featured spatial offsets relative to the sound of same magnitude (Δ) but opposing directions (arrows). In experiment 4 the two visual stimuli featured spatial offsets that were independent of each other and could have the same and opposing directions and distinct magnitudes. Experiment 2-4 also included auditory trials (A) in which only a sound was presented, and auditory trials were always preceded by an AV trial (for more details please refer to the main text). Sounds were presented from one of five discrete locations (indicated by magenta + signs in the A trial) and sound locations were pseudorandomized and counterbalanced across trials and conditions. Participants responded by moving a mouse cursor to indicate the perceived location of either a sound (in most trials) or a visual stimulus (see main text). The visual stimuli were presented with specific spatial discrepancies around each sound location and differed in color. The AV trials were used to probe the ventriloquism effect and to induce the aftereffect, the A trials were used to quantify the aftereffect.

Stimulus presentation was controlled using the Psychophysics toolbox (Brainard, 1997) for MATLAB (The MathWorks Inc., Natick, MA) with confirmed temporal synchronization of auditory and visual stimuli. The acoustic stimulus was a 1300 Hz sine wave tone (50 ms duration; sampled at 48 kHz) that was presented at 65 dB SPL through one of 5 speakers (Monacor MKS-26/SW, MONACOR International GmbH & Co. KG). These speakers were located behind the screen at 5 discrete horizontal locations (±22°,±11°, 0°), with the center position always being directly in front of the participant. Sound presentation was controlled via a multi-channel soundcard (Creative Sound Blaster Z) and amplified via an audio amplifier (t.amp E4-130, Thomann). The visual stimuli were projected (LG HF65, LG electronics) onto the acoustically transparent screen. The visual stimulus was a cloud of dots distributed according to a two-dimensional Gaussian distribution (200 dots, SD of vertical and horizontal spread = 1.6°, width of a single dot = 0.12°, duration 50 ms), similar to previous studies (Park & Kayser, 2019; 2021). The color of the dots differed between experiments. For experiment 1 these were white (rgb: 250, 250, 250), for experiments 2 to 4 these were orange or blue to allow their differentiation in the task instructions and to perceptually separate two visual stimuli presented in the same trial. The precise values of the blue (rgb 0, 65, 205) and orange hue (rgb 160, 60, 0) were chosen so they had similar perceptual saliency (luminance) when projected on the screen. These rgb values were obtained by letting five individuals adjust the overall luminance of the hues to equate their perceptual similarity. We did not probe participants for deficits in color vision, though we note that all participants were able to perform the task well and we did not observe systematic differences in the judgement errors between colors. The visual stimuli were either presented at the same discrete locations as the sounds or were positioned relative to the sound locations (see below). The individual locations of auditory and visual stimuli within trials (for AV trials) and between trials were drawn semi-independently resulting in stimulus positions that corresponded to different degrees of spatial (audio-visual) discrepancies.

Before each experiment participants were given the chance to familiarize themselves with the task and stimuli by performing 20 to 40 practice trials, for which they received feedback on their responses. The practice block included A, AV and V trials, similar to the main experiments. Before the actual experiment participants were also tested for their spatial hearing abilities. In a screening task, participants were asked to localize noise bursts (50 ms duration, 65 dB SPL) presented from the 4 lateralized speakers in a left/right two-choice task, performing 10 repeats per location. Participants performing below an average of 75% correct responses were not invited for the actual experiments (4 out of 95 tested individuals). The average performance of the included 91 participants was 93 % correct (mean; range 78% to 100%).

### Experiment 1

This experiment was designed to test how the stimulus onset asynchrony (SOA) between one visual stimulus and the sound influenced the ventriloquism effect. The SOAs covered five levels: ±600 ms, ±300 ms, and 0 ms. The individual auditory and visual stimuli were centered on one of five horizontal locations (±22°,±11°, 0°, the latter being straight ahead), with the auditory and visual stimuli always being spatially offset at discrepancies of ±33°, ±22°, and ±11° (defined as visual minus auditory stimulus position). Participants were asked to localize the sound in the AV trials, or to localize the visual stimulus in less frequent V trials. Each SOA was presented 84 times (14 times per spatial discrepancy) in pseudo-random order. For this experiment we collected data from 23 participants, of which one dataset had to be excluded (see below).

### Experiment 2

This experiment was designed to compare sound localization judgements in the presence of one or two visual stimuli. In the case of one visual stimulus this could be synchronous with the sound (condition AV0) or delayed by 300 ms (AV+300). In the case of two visual stimuli one was synchronous and one delayed (AV0V+300; Figure 1). The delay of 300 ms was determined based on the results from experiment 1. The auditory stimuli could appear at one of five horizontal locations (±22°,±11°, 0°) and the visual stimuli were placed systematically around these five sound locations, implementing six relative spatial discrepancies (±10°,±6°,±2°). In trials with two visual stimuli, one was displaced to the left of the sound and the other to the right. Hence, the two visual stimuli always featured opposing spatial discrepancies of same magnitude. During AV trials participants were asked to either localize the sound or one of the visual stimuli. To facilitate this, the two visual stimuli differed in color (blue or orange, with the assignment of V0 and V+300 to each color being pseudo-randomized across participants). Note that we here only analyze the data from trials requiring an auditory judgement. To test whether the influence of the delayed visual stimulus is modulated by the position of this relative to the sound (rather than just its presence) this experiment also included a control condition in which a delayed visual stimulus was presented at the same location as the sound and only the synchronous visual stimulus was spatially offset (AV0Vcontrol). Participants performed 960 trials, split across 10 blocks. These 960 trials reflect 480 AV trials (60 per each of the eight AV conditions defined by stimulus composition, 30 of which required an auditory judgment and 30 a visual judgment), each one followed by one of 480 A trials. For each condition the 60 trials were equally divided across the five sound locations and the six spatial discrepancies. Conditions were presented in pseudo-random order. We collected data from 21 participants, of which one had to be excluded.

### Experiment 3

This experiment extended the insights from experiment 2 to conditions in which the visual stimuli could appear either 300 ms prior to the sound (V-300), be synchronous with this (V0), or be delayed by 300 ms (V+300). As in experiment 2, the visual stimuli were placed around the sounds at six spatial discrepancies (±10°,±6°,±2°), and in trials featuring two visual stimuli, these had spatial offsets of same magnitude but opposing directions. In this experiment the vast majority of AV trials required participants to judge the sound, and visual judgements were required only in infrequent trials to maintain attention on both modalities. We reduced the fraction of visual judgements in this experiment as the results from experiment 2 revealed a minimal ventriloquism bias following visual judgements. Participants performed 936 trials in total, split across 12 blocks. These included 360 AV trials with an auditory judgement, 108 AV trials requiring participants to judge one of the visual stimuli, and 468 A trials, each one following an AV trial. The AV trials with an auditory judgement were equally divided across the five sound locations and the six spatial discrepancies. We collected data from 26 participants, of which two datasets had to be excluded.

### Experiment 4

This experiment was designed to extend the above insights to conditions in which two visual stimuli could feature distinct and independent spatial offsets to the sound. Similar to experiment 3 the visual stimuli could either precede the sound, be synchronous or delayed. However, in trials featuring two visual stimuli these could have spatial offsets in opposing directions but same magnitude ([+9,-9],[-9,+9], [+3,-3],[-3,+3]), opposing directions and distinct magnitudes ([+9,-3],[-3,+9],[-9,+3],[3,-9]), or could point in the same directions but have distinct magnitudes (([+9,+3,[+3,+9],[-9,-3],[-3,-9]). As a result, the spatial discrepancies of the two visual stimuli were independent of each other. The vast majority of AV trials required participants to judge the sound. Participants performed 900 trials each, split across 5 blocks. These included 360 AV trials with an auditory judgement, 360 A trials, and 180 AV trials requiring participants to judge one of the visual stimuli. We collected data from 21 participants.

### Definition of trial-wise judgement errors

For each AV and A trial we computed the deviation between the participant’s trial-wise response and the expected response given the location of the sound in that trial (Kording *et al*., 2007; Wozny & Shams, 2011a; b; Park & Kayser, 2019). This expected response captures potential response tendencies, such as to judge stimuli closer to the midline of the screen, but is not shaped by the ventriloquism bias given the overall balancing of stimulus positions and audio-visual discrepancies around each sound location. Practically, the expected response was computed for each sound location as the average response across all A trials with a sound at this location. The trial-wise judgement error was then defined as the actual response on that trial minus the expected response.

### Data screening

We removed few individual trials with judgement errors larger than 150% of the largest spatial discrepancy included in each experiment. These trials can reflect lapses of attention or accidental clicks when submitting the response. We visually inspected the resulting biases (as defined below) for each participant. This revealed very large response errors for 4 datasets (typically at least twice as large as the average error across all participants), which can for example arise when participants do not follow task instructions (e.g., localizing the visual rather than the auditory stimulus). These four participants were excluded from subsequent analysis, and the reported results are from n=22, 20, 24 and 21 datasets respectively for experiments 1 to 4. For those datasets retained the fraction of removed trials constitutes on average less than 0.5%.

### Analysis of the ventriloquism biases and aftereffects

The ventriloquism bias was obtained based on the AV trials requiring a judgement of sound location, while the aftereffect was obtained based on the A trials subsequent to these AV trials. The respective judgement errors were analyzed as a function of the audio-visual spatial discrepancy, defined as the location of the visual stimulus minus that of the sound in AV trials (Figures 2A,3A,B left, 4A). For conditions featuring one visual stimulus the relevant discrepancy is that of the one visual stimulus. For conditions featuring two visual stimuli, we proceeded as follows. In experiments 2 and 3 the visual stimuli featured directly opposing (hence anti-correlated) discrepancies. Hence, the bias expressed relative to one visual stimulus is the sign-inverted bias relative to the other and either discrepancy could be used as a reference (except for a difference in sign). In practice we used the visual stimulus yielding a positive slope (i.e. the one attracting the sound localization judgement). In experiment 4 the two visual stimuli featured independent discrepancies and hence both discrepancies need to be considered in the analysis. To analyze the data from all experiments in a comparable manner, we relied on participant-wise regression analyses to characterize the (linear) slope of the trial-wise judgement error against the spatial discrepancy of each visual stimulus (Debats *et al*., 2021; Park & Kayser, 2022). One regression was fit for each participant and condition. Both the dependent and independent variables were z-scored within each regression model to allow a comparison within and between conditions and experiments. Though we note that this z-scoring had essentially no influence on the overall conclusions. For conditions with one visual stimulus or conditions involving two anti-correlated visual stimuli the predictors were the spatial discrepancy of one visual stimulus and an offset; for conditions with two independent visual stimuli the predictors were both spatial discrepancies, their interaction, and an offset. In the last case the regression looked as follows, here exemplified for condition AV0V+300:

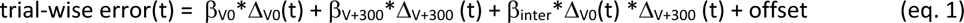

with Δ_V0_(t) denoting the trial-wise spatial discrepancy of the synchronous visual stimulus and Δ_V+300_(t) that of the delayed visual stimulus. The interaction term was included to allow a direct comparison of the influence of the two visual stimuli, though we verified that the main results do not depend on the inclusion of this interaction. The resulting standardized regression beta’s (termed slopes in the following) were considered as participant-wise measure of the respective contribution of each visual stimulus to the ventriloquism effect or aftereffect. These are shown in Figures 2B, 3A and B right, and 4B,C.

**Figure 2.**
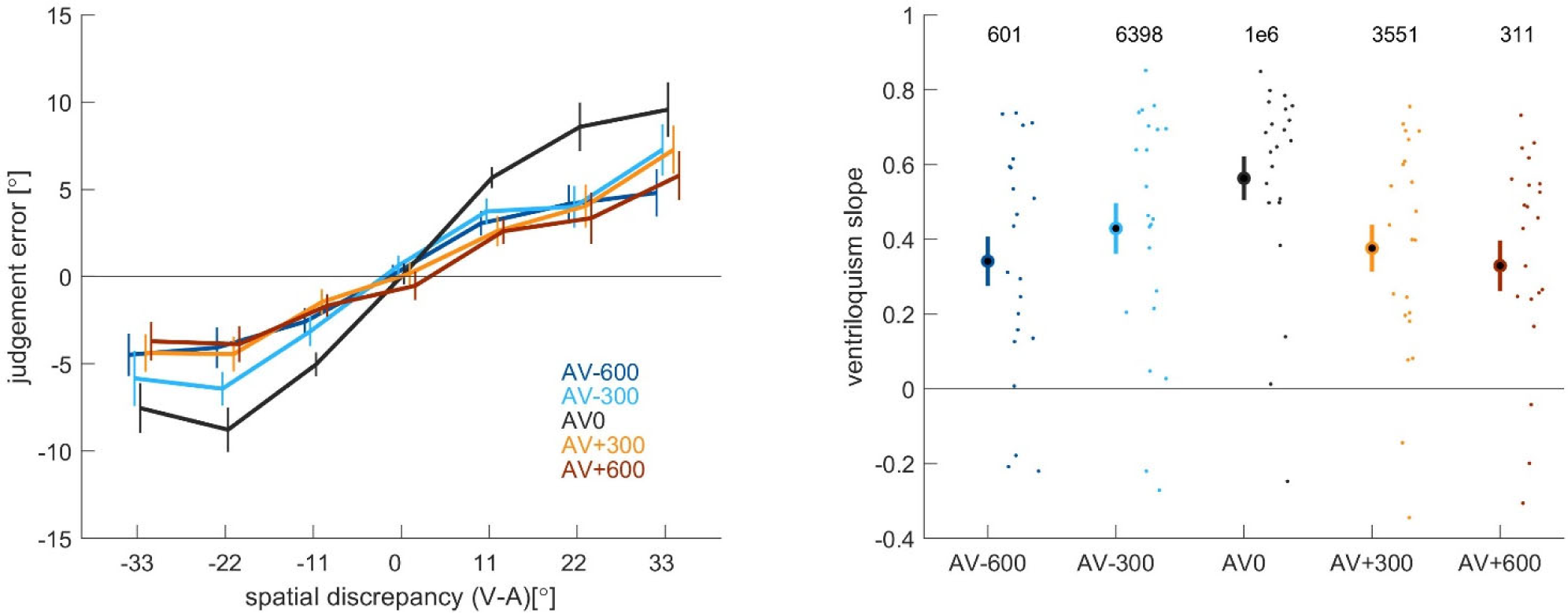
Dependency of the ventriloquism effect on SOA. Results from experiment 1 testing how the stimulus onset asynchrony (SOA) between one visual stimulus and a sound influences the ventriloquism effect. Left: Group-level judgement errors as a function of spatial discrepancy (defined as the position of the visual stimulus minus the position of the sound). Individual conditions are labeled by the SOA: AV-600 indicating that the visual stimulus precedes the sound by 600ms. Right: Regression slopes quantifying the ventriloquism effect. Numbers on top indicate the Bayes factor in favor of the group average to differ from zero (two-sided t-tests). Lines and circles indicate the group mean, error-bars the s.e.m. across participants, dots the individual participants (n=22).

**Figure 3.**
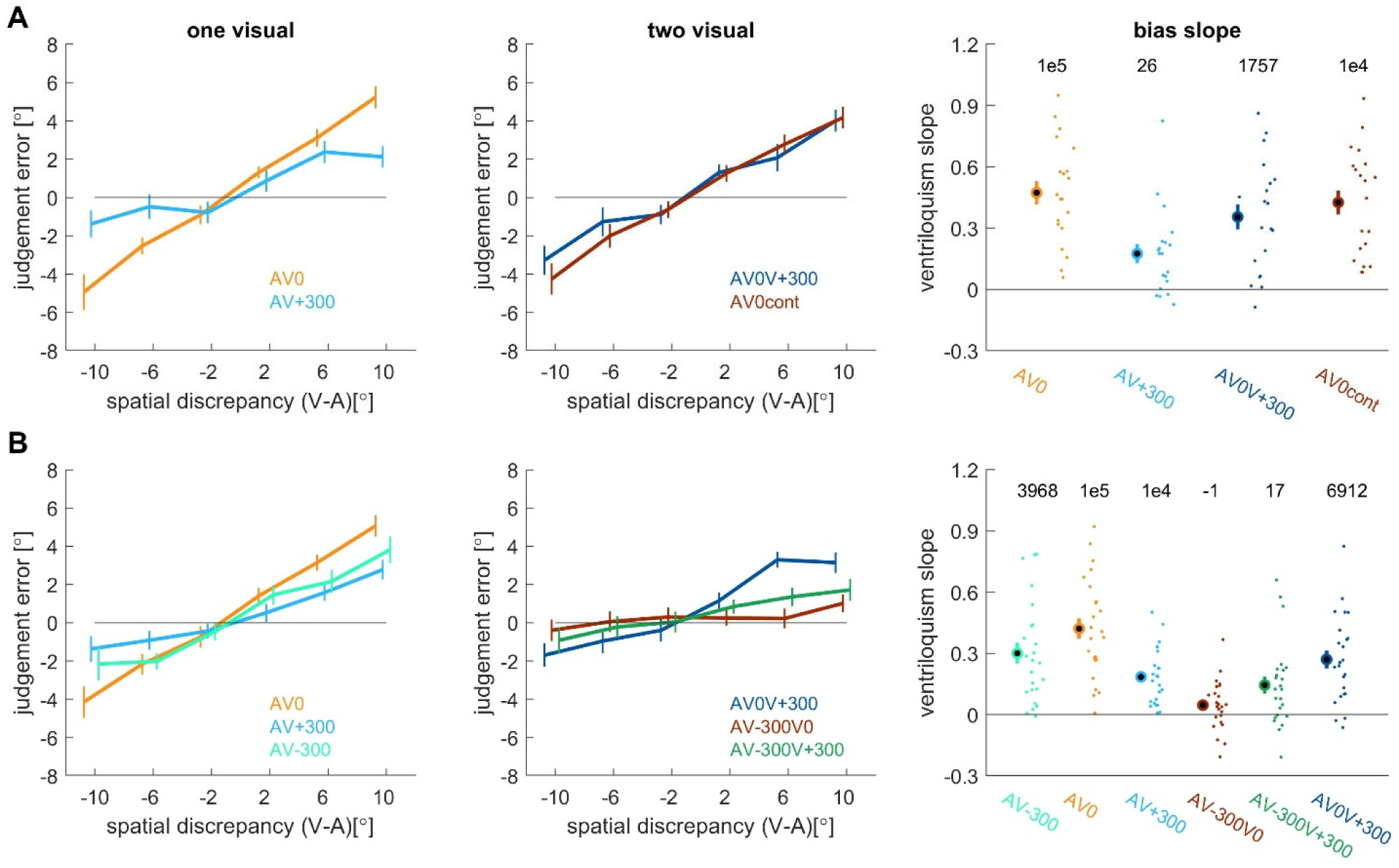
Ventriloquism bias for one and two visual stimuli (experiments 2 and 3). Left: Group-level judgement errors as a function of spatial discrepancy for AV trials with one visual stimulus. Middle: same for AV trials with two visual stimuli. Right: Slopes characterizing the ventriloquism effect. Numbers on top indicate the Bayes factor in favor of the group average to differ from zero (two-sided t-tests). **A**: Data from experiment 2 (n=20). **B**: Data from experiment 3 (n=24). Spatial discrepancy is defined as the position of the visual stimulus (when one was presented, or the one indicated first in the condition label) minus the position of the sound. Note that in these experiments the two visual stimuli presented in the same trial had opposing spatial discrepancies. For example, the vanishing bias for AV-300V0 in experiment 3 indicates a comparable influence of both visual stimuli. Lines and circles indicate the group mean, error-bars the s.e.m. across participants, dots the individual participants.

**Figure 4.**
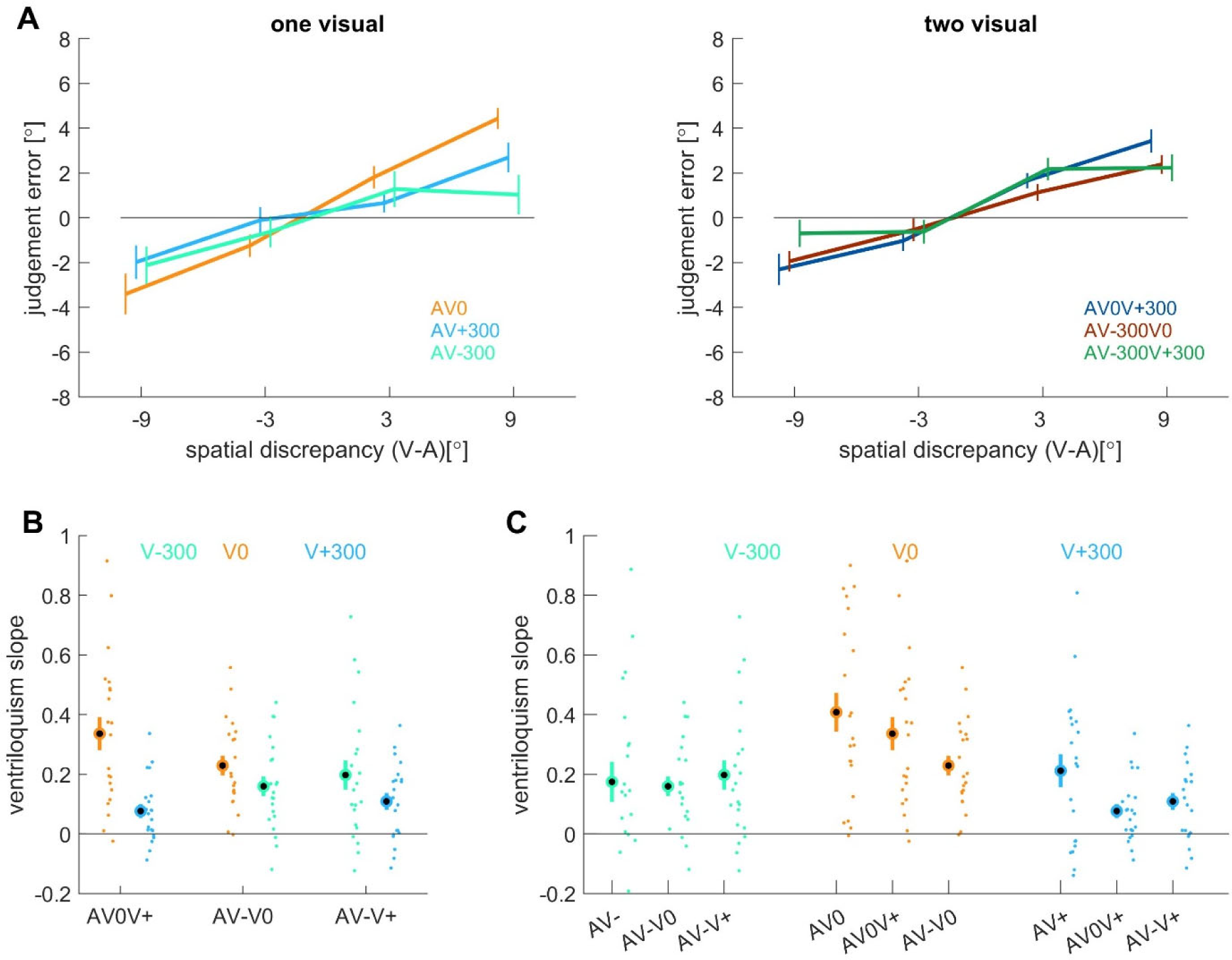
Ventriloquism bias for one and two visual stimuli (experiment 4). **A:** Group-level judgement errors as a function of spatial discrepancy for AV trials with one visual stimulus (left) and two visual stimuli (right). **B:** Regression slopes characterizing reflecting the contribution of the two visual stimuli in conditions featuring two visual stimuli (labelled on the x-axis). Their interaction terms were not significant (see main text). The colors reflect the SOA of individual stimuli. **C:** Systematic comparison of the regression slopes of visual stimuli at specific SOAs (color-coded) across conditions featuring one or two visual stimuli (labelled on the x-axis). This figure shows how the relative contribution of a visual stimulus prior to the sound (V-300) remains comparable across conditions, while that of a synchronous (V0) or a delayed (V+300) visual stimulus differ among conditions. Lines and circles indicate the group mean, error-bars the s.e.m. across participants, dots the individual participants. Some conditions are abbreviated (e.g. AV-denotes AV-300 and AV0V+ denotes AV0V+300).

The ventriloquism bias can in principle taper off at large discrepancies, reflecting the reduced binding of spatially very discrepant signals as predicted by models of Bayesian multisensory inference (Kording *et al*., 2007; Rohe & Noppeney, 2015; Shams & Beierholm, 2022). While this is seen in the paradigm used here when larger spatial discrepancies are included (Park & Kayser, 2020; Park *et al*., 2021; Park & Kayser, 2022), the present experiments were restricted to a smaller range of discrepancies and the resulting biases were largely linear. We verified this by comparing the quality of fit of a linear slope for the bias with a model featuring a linear plus a non-linear term, similar as we have used in previous studies (Cao *et al*., 2019; Park *et al*., 2021; Park & Kayser, 2022). This non-linear term was proportional to the square-root of the discrepancy and can describe the tapering off at very large spatial discrepancies. For AV trials with one visual stimulus in experiments 2 and 3 the average r-square fit improved only marginally from 0.130±0.020 to 0.138±0.019 (mean±s.e.m. of the participant wise qualities of fit averaged across AV conditions with a single visual stimulus).

### Analysis of error distributions

We probed whether the distribution of trial-wise judgement errors systematically differs between conditions featuring one or two visual stimuli. In particular we asked whether both are well accounted for by a unimodal distribution or whether they show signs of a potential switching of stimulus-response strategies and hence a bimodal distribution (Wozny *et al*., 2010; Rohe & Noppeney, 2015). To this end we computed the standard deviation (SD) of the judgement errors per condition and averaged these among conditions featuring one or two visual stimuli, per experiment and participant. If judgments following two visual stimuli are guided by a switching of decision strategies or multisensory computations (e.g. winner-takes-all in some trials and a weighted average in others), we would expect the SD to be larger for conditions featuring two visual stimuli.

### Statistical analysis and hypothesis testing

The main comparisons of interest concerned the group-level slopes reflecting the influence of the visual stimuli on sound localization judgements. To probe whether two visual stimuli exert a comparable influence, experiments 2 and 3 featured visual stimuli with directly opposing discrepancies. For these conditions we contrasted the slopes to a null hypothesis of zero, which reflects a direct cancellation of the two influences (Table 2). To probe the same hypothesis for experiment 4 we compared the slopes of the two visual stimuli per condition with each other by inspecting their interaction term and comparing the group-level standardized regression coefficients. We also probed whether the collective bias induced by two visual stimuli can be described by a capture-like model, in which one visual stimulus dominates the bias and the other contributes negligibly. In experiments 2 and 3 this would be reflected by a slope relative to one stimulus that is comparable to the bias induced by this visual stimulus when presented alone with the sound (Table 2). In experiment 4 such a scenario would be reflected by the weights of one visual stimulus to vanish. These tests were implemented using paired t-tests, for which we report t-values, p-values and Bayes factors (BF) against the null hypothesis (evidence in favor of the null is coded as negative values; hence transformed as - 1/BF). Bayes factors were obtained using the BayesFactor toolbox for Matlab (DOI:10.5281/zenodo.7006300). When interpreting Bayes factors (BF) we refer to the nomenclature of Raftery (Raftery, 1995) and interpreted BF between 1 and 3 as ‘weak’, between 3 and 20 as ‘positive’, between 20 and 150 as ‘strong’, and BF > 150 as ‘very strong’ evidence. Note that we base our interpretation of the statistical outcomes on the Bayes factors rather than the p-values in line with recommendations for behavioral studies (Wagenmakers, 2007; Wagenmakers *et al*., 2018). For some individual comparisons one could in principle have combined the data from all three experiments into a single linear model. However, we refrained from doing so for two reasons: first, the context of individual experiments can potentially induce experiment-specific factors that may be lost in such an analysis (Park & Kayser, 2022); and because the data were collected anonymously some participants may have participated in more than one of these experiments, which makes the handling of this factor in a combined analysis difficult.

We also probed an influence of the stimulus onset asynchrony on the bias observed for individual visual stimuli. For experiment 1 this was implemented using linear models, for the other experiments using paired t-tests.

## Results

### The ventriloquism effect follows a temporal binding window

In the first experiment we probed how the ventriloquism bias depends on the stimulus onset asynchrony (SOA) between the visual stimulus and the sound. We expected the ventriloquism bias to follow the general principle of a multisensory temporal binding window, hence to be strongest for synchronous stimuli (van Eijk *et al*., 2008; Diederich & Colonius, 2009; Stevenson & Wallace, 2013; Wallace & Stevenson, 2014). Indeed, the data (Figure 2) reveal a clear effect of SOA, and a linear model provided very strong evidence in favor of an effect of the magnitude of the SOA on the ventriloquism slope (contrasting 0, ±300, ±600: F(2,107)=38.3, p<10^-5^, BF∼10^8^, n=22; Figure 2). Subsequent paired comparisons provided positive evidence for no effect of SOA direction, hence comparable slopes for visual-leading and auditory-leading trials, and strong evidence for a reduced slope when the SOA magnitude becomes larger (Table 1). The data obtained in experiments 2,3 and 4 (the conditions featuring only one visual stimulus) further confirmed these findings (Table 1). Collectively this suggests that in the present experiments the ventriloquism bias induced by one visual stimulus decreases with SOA magnitude but shows little sensitivity to the specific order of the auditory and visual stimuli. Based on these results we selected an SOA of ±300 ms for the subsequent experiments, as it induces sufficient multisensory binding of visual and auditory stimuli, but at the same time assures that the bias induced by a synchronous and an asynchronous stimuli differ in strength.

**Table 1.**
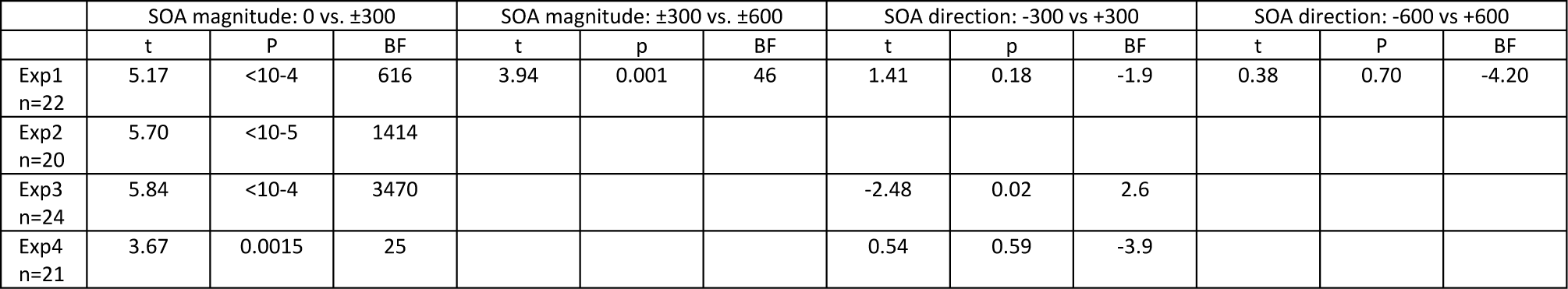
Dependency of the ventriloquism effect on SOA. Results from statistical comparisons (paired two-sided t-tests) of the group-level ventriloquism slopes between conditions. For conditions marked as ±, the slopes were averaged across the two conditions of same SOA magnitude by different sign. Empty cells reflect conditions not present in that experiment. Sample sizes are indicated (n). BF=Bayes factor, with positive values indicating evidence against the null hypothesis of no difference (between conditions or from zero).

| | SOA magnitude: 0 vs. $\pm 300$ | | | SOA magnitude: $\pm 300$ vs. $\pm 600$ | | | SOA direction: -300 vs +300 | | | SOA direction: -600 vs +600 | | |
| --- | --- | --- | --- | --- | --- | --- | --- | --- | --- | --- | --- | --- |
|  | t | P | BF | t | p | BF | t | p | BF | t | P | BF |
| Exp1<br>n=22 | 5.17 | <10<sup>-4</sup> | 616 | 3.94 | 0.001 | 46 | 1.41 | 0.18 | -1.9 | 0.38 | 0.70 | -4.20 |
| Exp2<br>n=20 | 5.70 | <10<sup>-5</sup> | 1414 |  |  |  |  |  |  |  |  |  |
| Exp3<br>n=24 | 5.84 | <10<sup>-4</sup> | 3470 |  |  |  | -2.48 | 0.02 | 2.6 |  |  |  |
| Exp4<br>n=21 | 3.67 | 0.0015 | 25 |  |  |  | 0.54 | 0.59 | -3.9 |  |  |  |

### The ventriloquism effect is shaped by both visual stimuli

In experiments 2 and 3 we probed whether two visual stimuli presented at different SOAs but with same spatial distance to the sound exert a comparable influence on the resulting ventriloquism bias. The respective judgement errors are shown as a function of the spatial discrepancy together in Figure 3. In the conditions featuring one visual stimulus synchronous with the sound and one delayed (or preceding the sound) the collective bias pointed towards the synchronous stimulus. In the condition involving two temporally offset stimuli (AV-300V+300) the bias pointed towards the visual stimulus preceding the sound (V-300). We reasoned that if both visual stimuli shape sound localization judgements in a comparable manner, one should observe a direct cancellation of their individual influences and hence a vanishing slope in the combined condition. In contrast, a differential influence of each visual stimulus should lead to a collective bias that differs from zero. In the extreme case of a winner-takes-all scenario the collective bias should be comparable to that observed in the condition featuring only the dominant visual stimulus. Table 2 summaries the statistical tests probing these possibilities.

**Table 2.** Analysis of ventriloquism slopes for experiments 2 and 3. The table reports comparisons of group-level slopes obtained with two visual stimuli to conditions involving only one visual stimulus or to a null effect of a mean slope of zero. Results are obtained from two-sided paired t-tests within experiments. Empty cells indicate conditions not present in the respective experiment. Sample sizes are indicated. BF=Bayes factor, with positive values indicating evidence against the null hypothesis of no difference (between conditions or from zero).

|  | AV0V+300 vs. AV0 |  |  | AV0V+300 vs. 0 |  |  | AV-300V0 vs. AV0 |  |  | AV-300V0 vs. 0 |  |  | AV-300V+300 vs. AV-300 |  |  | AV-300AV+300 vs. 0 |  |  |
| --- | --- | --- | --- | --- | --- | --- | --- | --- | --- | --- | --- | --- | --- | --- | --- | --- | --- | --- |
|  | T | P | BF | t | p | BF | T | p | BF | t | P | BF | t | p | BF | T | p | BF |
| Exp2<br>n=20 | -3.58 | 0.002 | 20 | 5.83 | <10<sup>-4</sup> | 1756 |  |  |  |  |  |  |  |  |  |  |  |  |
| Exp3<br>n=24 | -4.32 | 0.0002 | 118 | 6.15 | <10<sup>-5</sup> | 6912 | -6.71 | <10<sup>-5</sup> | 23049 | 1.84 | 0.078 | -1.1 | -4.97 | <10<sup>-4</sup> | 503 | 3.47 | 0.002 | 17 |

The collective data speak in favor of a differential influence of each visual stimulus and against the dominance of a single stimulus. Specifically, when one visual stimulus is synchronous with the sound and one delayed (AV0V+300) the collective bias differs from zero providing strong support for a differential contribution of both visual stimuli (Table 2; Figure 3B). The same result was obtained when both visual stimuli were temporally offset from the sound (AV-300V+300). Only when one visual stimulus preceded the sound and one was synchronous (AV-300V0) did we obtain no clear evidence for a difference from zero. When comparing the collective bias against that induced by the strongest individual visual stimulus, we obtained very strong evidence for the collective bias to be smaller (Table 2).

To further characterize the combined influence of two visual stimuli, experiment 2 featured a condition in which one visual stimulus was synchronous to and spatially offset from the sound and a second visual stimulus (delayed by 300 ms) was presented at the same location as the sound (condition AV0Vcontrol). The rationale was the following: if the influence of this delayed visual stimulus emerges only in direct proportion to its spatial discrepancy, it should not contribute to the collective bias in the control condition (as the discrepancy is zero). Hence the collective bias in this control condition should be comparable to that obtained for only the synchronous visual stimulus. Indeed, the data show weak evidence in support of this (AV0Vcontrol vs. AV0, t=-1.85, p=0.079, BF=-1.0) as the overall bias was smaller (Figure 3A). At the same time the bias in the control condition also differed from that obtained when the delayed visual stimulus was spatially offset from the sound (AV0Vcontrol vs. AV0V+300; t=2.83, p=0.01, BF=4.9). This suggests that a second visual stimulus shapes the bias not only in direct proportion to its spatial offset but also by adding to the overall visual context present in each trial.

In experiment 4 the two visual stimuli featured independent spatial offsets that could vary in magnitude and direction (see Figure 4A for judgement errors). In this experiment, directly opposing influences of the two visual stimuli do not necessarily lead to a vanishing bias in the combined condition. To determine whether two visual stimuli in a given condition had a comparable influence we compared their individual slopes and quantified their interaction (Figure 4B). We obtained positive evidence was positive for the interaction to be negligible (t-values between -0.79 and 0.45, BF between -3.33 and -4.0). We hence compared the standardized slopes between the two visual stimuli within and between conditions.

The contrast within conditions provided strong evidence for a differential influence of a synchronous and a delayed visual stimulus when combined (slopes within AV0V+300: t=4.01, p=0.0004, BF=74) and no clear result for the other conditions (AV-300V0: t=2.05 p=0.053, BF=1.3; AV-300V+300: t=1.67, p=0.11, BF=-1.3). We then further compared the contribution of a visual stimulus at specific SOAs between all conditions, those featuring one or two visual stimuli (Figure 4C; Table 3). This provided positive evidence for the influence of a visual stimulus prior to the sound (V-300) to be comparable regardless of whether it is presented without a second visual stimulus or not (Table 3). In contrast, this provided weak to positive evidence for the influence of a synchronous visual stimulus to be strongest when present in isolation. For a delayed visual stimulus (V+300) this provided weak to positive evidence for the bias to be stronger when was presented without a second visual stimulus and weak evidence against a difference when combined with another visual stimulus.

**Table 3.**
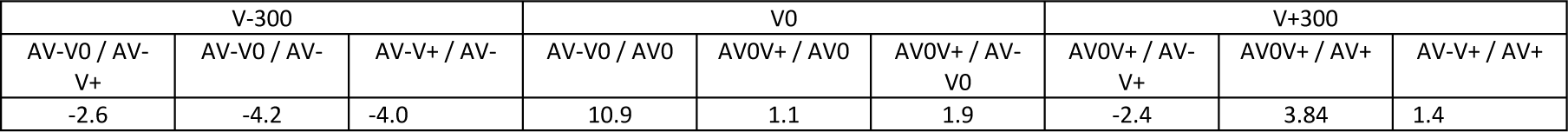
Comparison of the slopes of visual stimuli at specific SOAs between conditions (experiment 4). The table reports the Bayes factors for comparisons of the group-level slopes obtained in different conditions (two-sided paired t-tests). Positive values indicate evidence against the null hypothesis of no difference between conditions.

| V-300 |  |  | V0 |  |  | V+300 |  |  |
| --- | --- | --- | --- | --- | --- | --- | --- | --- |
| AV-V0 / AV-V+ | AV-V0 / AV- | AV-V+ / AV- | AV-V0 / AV0 | AV0V+ / AV0 | AV0V+ / AV-V0 | AV0V+ / AV-V+ | AV0V+ / AV+ | AV-V+ / AV+ |
| -2.6 | -4.2 | -4.0 | 10.9 | 1.1 | 1.9 | -2.4 | 3.84 | 1.4 |

### Analysis of trial-wise response distributions

In models of multisensory inference the sensory evidence obtained under different assumptions about a causal relation between sensory signals can be combined for a perceptual response according to different decision strategies (Wozny *et al*., 2010; Rohe & Noppeney, 2015; Park & Kayser, 2022). Strategies that incorporate a binary choice between two options can lead to multimodal response distributions, whereby some trials are guided by one strategy and other trials by another. In the present experiment it could for example be that judgements in trials with two visual stimuli are sometimes guided by only one and sometimes by only the other of these and sometimes by a weighted combination of both. We implemented additional data analyses to probe whether the trial-wise response distributions in conditions featuring two visual stimuli are shaped differently than those in conditions with a single visual stimulus.

To this end we calculated the standard deviation of the participant- and condition-wise judgement errors (Figure 5). If multisensory combination for two visual stimuli is guided by some form of alternation between two decision strategies, we expected the standard deviations to be larger in those trials. However, the data suggest that this is not the case as the standard deviations were generally smaller in conditions featuring two visual stimuli (Experiments 2,3,4: t=3.32, p=0.003, BF=12; t=1.45, p=0.16, BF=-1.9; t=4.22, p=0.0004, BF=77). Hence, the trial-wise data do not provide evidence for a switching in response strategies between conditions featuring one or two visual stimuli.

**Figure 5.**
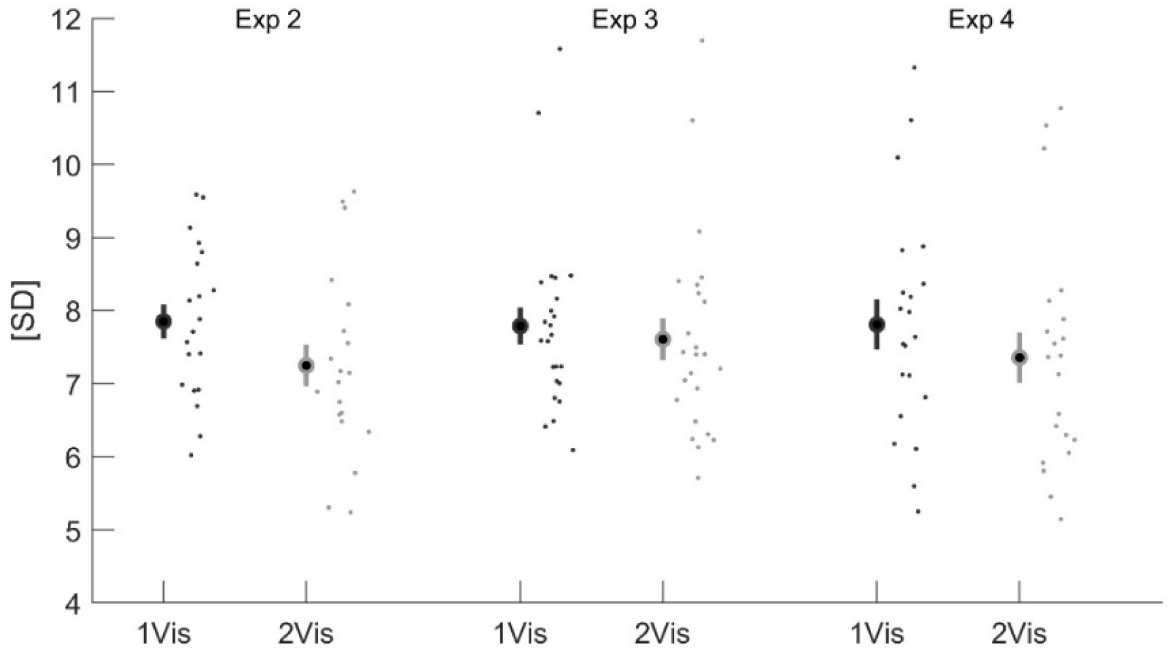
Distribution of trial-wise judgement errors. The graph shows the condition- and participant-wise standard deviations of the trial-wise judgement errors for each experiment. The standard deviation was computed for each condition and then averaged across conditions featuring one (1Vis) or two visual stimuli (2Vis). Circles indicate the group mean, error-bars the s.e.m. across participants, dots the individual participants.

### The ventriloquism aftereffect diminishes following two visual stimuli

The trial-wise ventriloquism aftereffect is a weak but robust bias visible in sound localization judgements following a single audio-visual trial. The present experiments allowed us to probe us how this aftereffect is shaped by the relative timing of the audio-visual stimuli and whether it emerges following AV trials featuring two visual stimuli. The respective judgement errors and regression slopes are shown in Figure 6. The aftereffects were robust following audio-visual trials featuring one visual stimulus but were weaker following those with two visual stimuli (see BF in Figure 6B). This was in particular the case for experiment 3 where the visual stimuli could precede or follow the sound (Table 4). However, contrasting the aftereffect slopes between conditions featuring one and two visual stimuli also yielded unclear results, making the specific interpretation of individual comparisons difficult. Still, the picture seems to be similar compared to the ventriloquism effect in that the collective visual evidence available in an audio-visual trial determines the aftereffect seen in a subsequent unisensory test trial.

**Figure 6.**
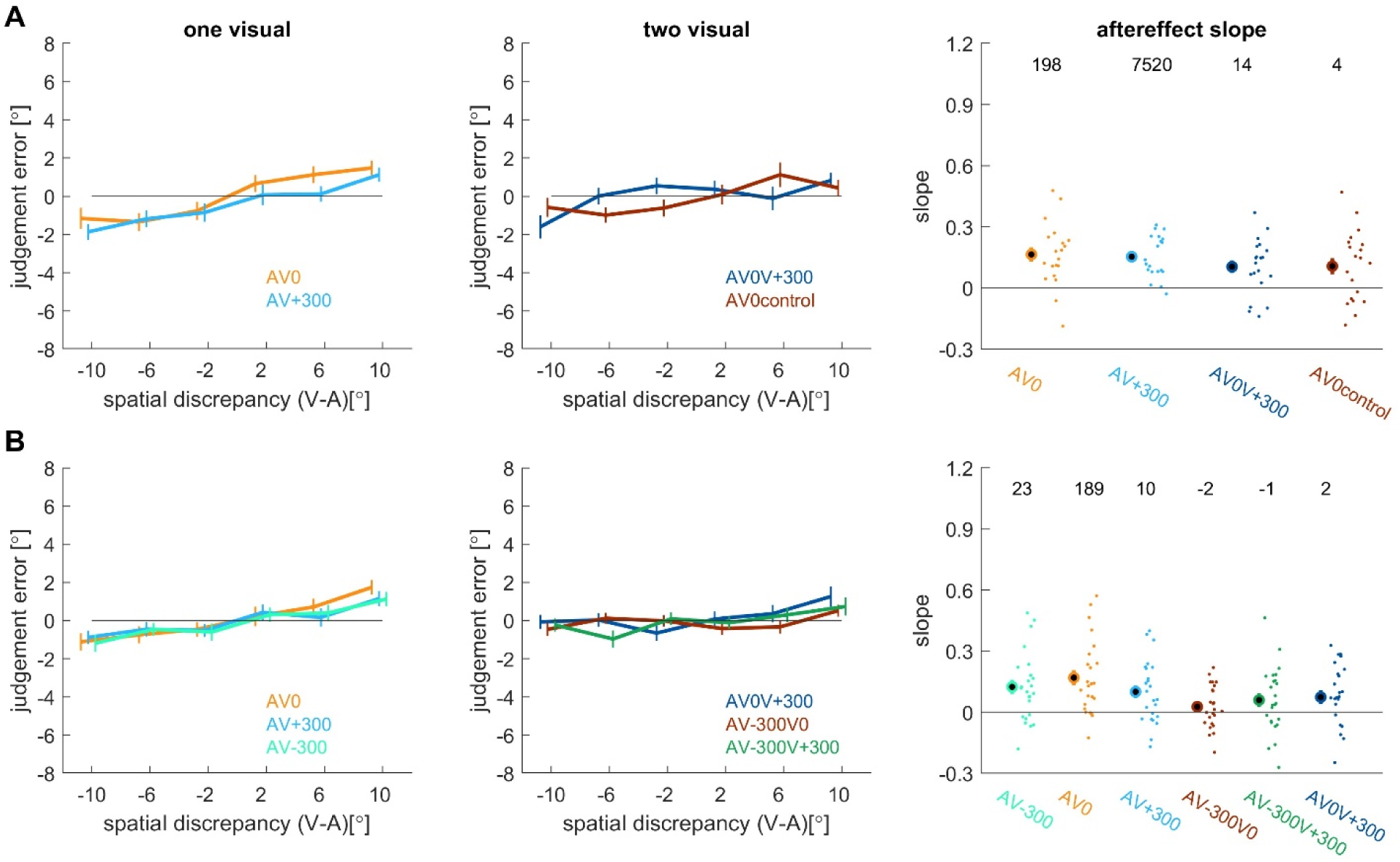
Ventriloquism aftereffect for one and two visual stimuli (experiments 2 and 3). Left: Group-level judgement errors as a function of spatial discrepancy for AV trials with one visual stimulus. Middle: same for AV trials with two visual stimuli. Right: Bias slope for each condition. Numbers on top indicate the Bayes factor for t-test contrasting the group average to zero. **A**: Data from experiment 2 (n=20). **B**: Data from experiment 3 (n=24). In the right panel conditions are abbreviated (e.g. AV-denotes AV-300 and AV0V+ denotes AV0V+300). Lines and circles indicate the group mean, error-bars the s.e.m. across participants, dots the individual participants.

**Table 4.** Analysis of aftereffect slopes for experiments 2 and 3. The table reports comparisons of group-level slopes obtained with two visual stimuli to conditions involving only one visual stimulus or to a null effect of a mean slope of zero. Results are obtained from two-sided paired t-tests within experiments. Empty cells indicate conditions not present in the respective experiment. Sample sizes are indicated. BF=Bayes factor, with positive values indicating evidence against the null hypothesis of no difference (between conditions or from zero).

|  | AV0V+300 vs. AV0 |  |  | AV0V+300 vs. 0 |  |  | AV-300V0 vs. AV0 |  |  | AV-300V0 vs. 0 |  |  | AV-300V+300 vs. AV-300 |  |  | AV-300V+300 vs. 0 |  |  |
| --- | --- | --- | --- | --- | --- | --- | --- | --- | --- | --- | --- | --- | --- | --- | --- | --- | --- | --- |
|  | t | p | BF | t | p | BF | t | p | BF | t | p | BF | t | p | BF | t | p | BF |
| Exp2<br>n=20 | -1.69 | 0.1 | -1.3 | 3.39 | 0.003 | 14 |  |  |  |  |  |  |  |  |  |  |  |  |
| Exp3<br>n=24 | -2.19 | 0.04 | 1.6 | 2.17 | 0.04 | 1.5 | -3.46 | 0.002 | 18.3 | 1.23 | 0.23 | -2.4 | -1.74 | 0.09 | -1.3 | 1.9 | 0.08 | -1.2 |

## Discussion

We asked how the presence of two visual stimuli shapes multisensory perception in the audio-visual spatial ventriloquism paradigm. Our results suggest that the biases in sound localization judgements arising from two visual stimuli are not only determined by the properties of each visual stimulus in relation to the sound alone, but by the collective configurational properties of all the available visual information. This result was clearly observed for the ventriloquism bias, but in our interpretation is likely to also hold for the immediate ventriloquism aftereffect. Depending on the overall configuration of the visual information, two visual signals can influence the collective spatial bias with comparable or with asymmetric weights that are shaped both by the temporal offset of each visual stimulus relative to the sound and by the temporal offset of the second visual stimulus.

### Sensory context shapes the multi-stimulus ventriloquism effect

Our results show that when two visual stimuli are combined with a sound both visual signals exert an influence on the ventriloquism bias. This speaks against the dominance, or capturing, of multisensory binding by a single multisensory stimulus pair. The effect of each visual stimulus depends on its own spatial and temporal distances to the sound, as is well known (Bertelson *et al*., 2000; Rohe & Noppeney, 2015; Bruns, 2019). However, our data show that the collective bias induced by two visual stimuli is not simply a direct sum of the biases seen when each visual stimulus is present in isolation. Rather, the collective effect is shaped by the overall configuration of the available visual information, rendering the outcome of multi-stimulus multisensory integration scene- or context-dependent. This result was consistently observed across three experiments that varied in the temporal onset asynchronies between the visual stimuli and the sound and the spatial discrepancies of both visual signals to the sound. Importantly, in experiment 4 the number of visual stimuli, their timing and their spatial offsets were unpredictable, suggesting that this result is not contingent on a specific expectation about the upcoming stimulus.

This context-sensitivity of the ventriloquism bias is best illustrated in Figure 4. This shows that the relative influence of visual stimuli that are synchronous or delayed to the sound differs depending on whether these are the sole visual stimulus or accompanied by another one. In contrast, the relative influence of a visual stimulus prior to the sound is constant across conditions. We interpret this context-sensitivity as a consequence of sensory inference shaping the binding process based on the collective sensory scene. Multisensory inference guides perception by fostering the binding of those sensory cues that are deemed to be causally related and to originate from a common sensory object (Kording *et al*., 2007; McGovern *et al*., 2016; Odegaard & Shams, 2016; Odegaard *et al*., 2017; Noppeney, 2021; Shams & Beierholm, 2022). This process requires establishing putative relations between sensory stimuli based on the spatio-temporal correlations between them, at least in the present experiments where other object-specific information was not present. During the course of an individual trial the timing and number of the visual signals was unpredictable. Hence, the first occurrence of any visual signal provides initial (and possibly sole) evidence for a multisensory correlation. We propose that once such evidence for a correlation, and hence a putative common cause, has been received subsequent signals receive a reduced weight, given that some evidence for a common relation has already been received. More specifically, the evidence for a causal relation of a visual stimulus presented after a sound is likely to be lower when this visual stimulus was preceded also by another visual stimulus at e.g. 50 ms prior to the sound. The temporally asymmetric emergence of evidence for putative causal relations results in an asymmetric contribution of visual signals to the collective bias. This notion predicts a comparable influence of a preceding and synchronous visual stimulus when only these are present, a reduced influence of a delayed one when paired with any other visual stimulus, and in absolute terms a stronger influence of a synchronous visual stimulus when presented as first visual stimulus. While the data obtained here directly supports such a model, the specific computational details of such a multi-stimulus multisensory causal inference model need to be worked out in detail and are beyond the scope of this more explorative study.

Interestingly, the differential influence of visual signals prior to and subsequent to a sound that we observed in conditions featuring two visual stimuli was not present in conditions with only one visual stimulus. Both in experiment 1, which featured only individual visual stimuli, and in the trials featuring one visual stimulus in experiments 2-4, was the ventriloquism bias for single visual stimuli independent of the temporal order of auditory and visual stimuli (sign of the SOA). This discrepancy between the temporal binding pattern observed for individual and pairs of visual stimuli in our view further substantiates the context-sensitivity of the ventriloquism effect. In particular, the precise SOA between sound and a visual signal may be more relevant for determining putative stimulus-relations when multiple rather than a single visual signal is present (Van der Burg *et al*., 2008a; Talsma *et al*., 2010), as in the latter case additional selection is required to determine which of two visual signals better relates to the sound (Kording *et al*., 2007; Noppeney, 2021; Shams & Beierholm, 2022). Hence the prevalence of temporal asymmetries in conditions featuring two visual signals may not be a qualitative difference to the more symmetric temporal pattern seen for individual visual stimuli here, but may simply be the consequence of the elaborate computations shaping sensory causal inference in more complex multisensory environments.

The present analysis was constrained to biases that were linearly proportional to the degree of spatial discrepancy and which combined the influence of temporally dispersed visual stimuli in a linear manner. For larger spatial separations the ventriloquism can scale nonlinearly with the spatial discrepancy (Kording *et al*., 2007; Rohe & Noppeney, 2015; Shams & Beierholm, 2022) and similarly, the temporal binding window can reflect a nonlinear interaction of multiple modalities at longer SOAs (Stevenson & Wallace, 2013). This poses limitations over the spatial range and time lags over which the present results may generalize and again calls for more detailed computational models of how multiple signals are combined within and across modalities.

We also note that the relative contribution of individual stimuli in real-life will depend on additional factors beyond their spatio-temporal alignment. It is known that the relative weight among two stimuli is shaped by task-relevance, semantic congruency, prior expectations and attention (Talsma *et al*., 2010; Tovar *et al*., 2020; Zuanazzi & Noppeney, 2020; Delong & Noppeney, 2021; Noppeney, 2021). In particular the role of attention is difficult to dissociate from other aspects of multisensory computation, as attention may be assigned unconsciously and automatically based on multiple attributes, including the timing of individual stimuli (Van der Burg *et al*., 2008b; Talsma *et al*., 2010; Donohue *et al*., 2015; Zhaoping, 2023). With that in mind, it is possible that attentional selection may contribute to the asymmetric contribution of visual stimuli prior to and subsequent to the sound in the ventriloquism bias, as observed here, and extreme forms of attention capturing may essentially look like multisensory capture.

### Aftereffect emerges robustly across task relevance and temporal delays

Similar to previous studies we observed a robust trial-wise (also called immediate) ventriloquism aftereffect in unisensory auditory trials following the audio-visual trials (Radeau & Bertelson, 1974; Recanzone, 1998; Bruns *et al*., 2011; Park & Kayser, 2019; 2021). For multisensory trials featuring one visual stimulus this aftereffect reflects a bias in the localization of a sound towards the direction of the spatial discrepancy experienced in the preceding multisensory trial (Wozny & Shams, 2011b; Park *et al*., 2016). Our results corroborate the robustness of this aftereffect and show that it emerges also when the visual and auditory stimuli are asynchronous, at least on the scale of 300 ms as used here. This fits the notion that the aftereffect emerges automatically, possibly as a result of the belief in a modality specific bias (Wozny & Shams, 2011a; Noppeney, 2021; Park & Kayser, 2021).

However, our data also show that following more ambiguous multisensory scenes, such as those including two visual stimuli, the aftereffect largely vanishes. In our view this result supports the notion that the aftereffect reflects an adjustment in response to a belief in a modality specific (i.e. auditory) bias. Potential evidence for such a bias emerges in the presence of individual audio-visual stimulus pairs, regardless of which stimulus was directly task relevant. However, when multiple visual signals are present it may be less clear whether or which one provides evidence about a modality specific bias. When performing a task based on an ambiguous or cluttered visual environment there may be no evidence for a modality specific bias in the auditory domain. This influence of task- or behavioral-relevance on the aftereffect is also supported by a study that rendered one of two visual stimuli behaviorally relevant over the course of an entire experiment and found an aftereffect bias specifically towards the relevant visual stimulus (Tong *et al*., 2020).

## Conclusion

In everyday life we are faced with multiple signals in each sensory modality and combining these into a coherent percept is a major feat achieved by the brain. While many laboratory studies employ only a single stimulus per modality, the present experiments pave the way to understand how multisensory perception behaves under more realistic conditions involving multiple stimuli in one modality. We suggest that the underlying sensory computations are guided by multisensory inference and the present data show how this results in context-sensitive multisensory biases in the spatial ventriloquism paradigm. Here, the influence of individual visual signals is shaped by their individual properties such as the spatial distance and temporal delay to a sound, but also by the temporal context provided by other visual signals. These results highlight the context-sensitive nature of multisensory perception, which so far has received comparatively little attention.

## Acknowledgements

This study was funded by the Deutsche Forschungsgemeinschaft (DFG KA2661/2-1). The authors declare that they have no conflicts of interest. We thank Jenny Jakisch and Lisa Stetza for their help during data collection.

